# Polyploidy of semi-cloned embryos generated from parthenogenetic haploid embryonic stem cells

**DOI:** 10.1101/2020.04.29.067991

**Authors:** Eishi Aizawa, Charles-Etienne Dumeau, Remo Freimann, Giulio Di Minin, Anton Wutz

**Affiliations:** Institute of Molecular Health Sciences, Swiss Federal Institute of Technology, ETH Zurich, Zurich, Switzerland

## Abstract

In mammals, the fusion of two gametes, an oocyte and a spermatozoon, during fertilization forms a totipotent zygote. There has been no reported case of natural parthenogenesis, in which embryos develop from unfertilized oocytes. The genome and epigenetic information of haploid gametes are crucial for the proper development of embryos. Haploid embryonic stem cells (haESCs) are unique stem cells established from uniparental blastocysts and possess only one set of chromosomes. Previous studies have shown that sperm or oocyte genome can be replaced by haESCs with or without manipulation of genomic imprinting for generation of mice. Recently, these remarkable semi-cloning methods have been applied for screening of key factors of mouse embryonic development. While haESCs have been applied as substitute of gametic genome, the fundamental mechanism how haESCs contribute to the genome of totipotent embryos is unclear. Here, we show the generation of fertile semi-cloned mice by injection of parthenogenetic haESCs (phaESCs) into oocytes after deletions of two differentially methylated regions (DMRs), the *IG*-DMR and *H19*-DMR. For characterizing the genome of semi-cloned embryos further we establish ESC lines from semi-cloned blastocysts. We report that polyploid karyotypes are observed frequently in semi-cloned ESCs (scESCs). Our results confirm that mitotically arrested phaESCs provide high efficiency for semi-cloning when the *IG*-DMR and *H19*-DMR are deleted. In addition, we highlight the occurrence of polyploidy that needs to be considered for further improvement for development of semi-cloned embryos derived by haESC injection.

## Introduction

The genetic information of an oocyte and a spermatozoon are inherited to the offspring. Both maternal and paternal genomic information are required for normal development of mammalian embryos because uniparental embryos cause developmental defect due to the imbalance of genomic imprinting [1]. Despite the importance of gametic genome, much remains to be elucidated what is required in genomic information of gametes to form a totipotent zygote. In mice, previous studies have shown that two differentially methylated regions within the *H19-Igf2* and *Gtl2-Dlk1* imprinted gene clusters had critical contributions to the genome of embryos [2, 3]. Both regions are normally methylated and unmethylated on the paternally and maternally inherited chromosomes, respectively. Deletion of the *H19*-DMR and the intergenic germline-derived DMR (*IG*-DMR) resulted in loss of expression of the maternally expressed *H19* and *Gtl2* genes from the maternal allele, respectively [4, 5]. Deletion of the *H19*-DMR or combined deletions of the *H19*-DMR and *IG*-DMR from the genome of non-growing oocytes facilitated the generation of bimaternal mice by substituting the manipulated genome of non-growing oocytes for sperm by injection into mature oocytes [2, 3].

Recent studies have also explored the possibility to substitute the gametic genome by mouse haploid embryonic stem cells (haESCs). Haploid ESCs are unique stem cell lines established from either parthenogenetic [6, 7] or androgenetic haploid blastocysts [8, 9]. Haploid ESCs possess a single set of chromosomes, that is 20 chromosomes in mice, similar to gametes. Due to their uniqueness of haploidy, haESCs have been applied to various studies in original manners. One application is for genetic screening. While heterozygous mutations in diploid cells are often masked phenotypically, hemizygous mutations in haploid cells directly express phenotypes. For example, gene trap vectors have been applied to screen genes required for chemical toxicity, self-renewal of ESCs and X-chromosome inactivation [6, 10-12]. Another considerable application of haESCs is as gametic genome. Several reports have demonstrated the application of either androgenetic haESC (ahaESC) or parthenogenetic haESC (phaESC) for substituting the paternally derived sperm genome to generate semi-cloned mice [8, 9, 13, 14]. Furthermore, it has been demonstrated that the oocyte genome can be replaced by the genome of phaESCs for generation of semi-cloned mice, albeit at low frequency [15]. In contrast to oocytes and spermatozoa, genetic mutations can be readily introduced to haESCs owing to their self-renewal capacity in culture. Methods for introducing genetic modifications into the germline with haESCs is a considerable approach for studies including embryonic development and generation of transgenic animals. Recent studies have already applied the combination of CRISPR-Cas9-based genome editing and haESCs as substitute of the gametic genome for genetic screening approaches [16], characterization of imprinting regions for embryonic development [17], and identification of important amino acids within the DND1 protein for primordial germ cell development [18].

While these remarkable studies have successfully applied haESCs as substitutes for gametic genomes, the mechanism how haESCs contribute to the genome of totipotent embryos remains to be clarified. For example, sperm and haESCs genomes are fundamentally different as most of the sperm genome is packaged with protamines, but the chromosomes of haESCs have a conventional nucleosomal structure. Proper segregation of both maternal and paternal haploid chromosome sets into each blastomere is required at the first division of the zygote to form a developmentally competent 2-cell embryo. Otherwise, developmental defects due to aneuploidy or polyploidy arise in embryos [19]. Polyploidy is the cell state possessing more than two complete sets of chromosomes, which is observed in various species including plants and yeasts [20]. In mammals, polyploid embryos can occur by polyspermy or abnormal chromosome segregation, but show developmental defects and arrest [20-22].

In this study, we report that the generation of healthy mice by injection of mitotically arrested phaESCs with manipulation of genomic imprinting into metaphase II (MII) oocytes. We established semi-cloned ESCs (scESCs) derived from semi-cloned blastocysts for characterization of their ploidy status. We find that scESCs exhibited polyploidy, indicating cautious analysis is required for the study of semi-cloned embryos generated by application of haESCs.

## Results

### Deletions of the *IG*-DMR and *H19*-DMR in haESC lines

A previous study has reported that bimaternal embryos generaterd by substituting the paternal genome of sperm by the haploid genome of non-growing oocytes cause developmental defects and arrest in embryogenesis [23]. These defects were largely overcome by manipulation of genomic imprinting by deleting the *IG*-DMR and *H19*-DMR from the genome of non-growing oocytes resulting in the development of bimaternal mice [2, 3]. These studies indicate that imprinted gene expression regulated by the *IG*-DMR and *H19*-DMR is the key barrier, which prevents the development in bimaternal embryos. In order to manipulate genomic imprinting in phaESCs, the CRISPR-Cas9 system was used to delete the *IG*-DMR and *H19*-DMR in a phaESC line that was established from a 129S6/SvEvTac mouse oocyte (Fig 1A and S1A-B Fig). After the culture of single cells transfected simultaneously with CRISPR-Cas9 plasmids and *piggyBac* transposon plasmids for EGFP expression, 2 phaESC lines, termed double-knockout phaESC line 1 (DKO-phaESC-1) and line 2 (DKO-phaESC-2), were selected by PCR-based genotyping (S1C Fig). DKO-phaESC lines were labelled with EGFP to distinguish embryos derived from DKO-phaESCs for further studies. DNA sequencing confirmed the deletions of 4,168 base pairs (bp) in the *IG*-DMRs for both DKO-phaESC-1 and DKO-phaESC-2 (Fig 1B). Also, deletions of 3,908 and 3,927 bp in the *H19*-DMRs were confirmed in DKO-phaESC-1 and DKO-phaESC-2, respectively. Both ESC lines had a similar morphology to the parental phaESC line (Fig 1C), and an intact haploid karyotype was confirmed by analysis of metaphase chromosome spreads (Fig 1D).

**Fig 1.**
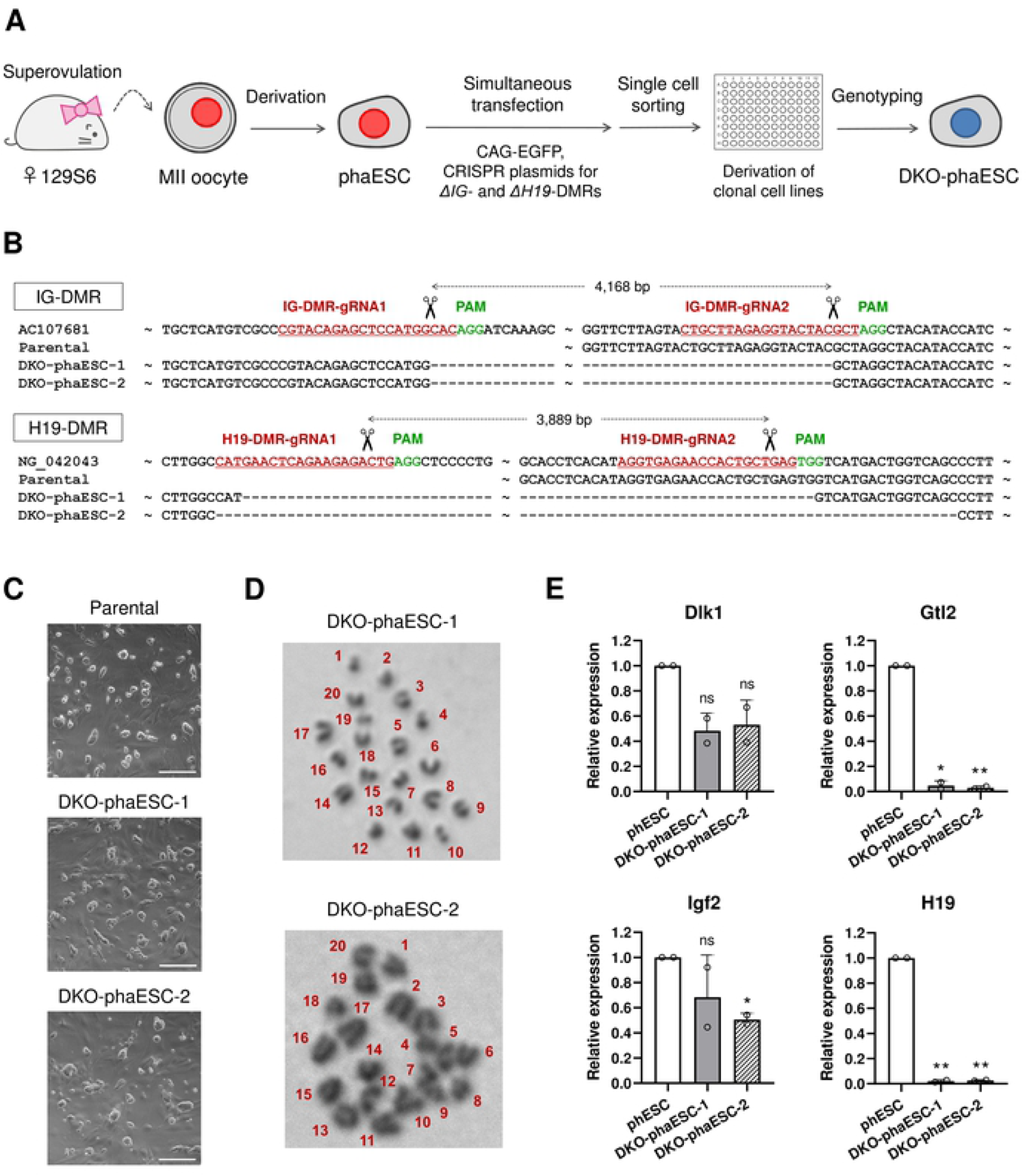
Generation of the *IG*-DMR and *H19*-DMR deletions in haESCs. (A) Strategy for generating the *IG*-DMR and *H19*-DMR deletions in phaESC lines that were established from activated mouse oocytes and were marked by integration of CAG-EGFP-IRES-hygro *piggyBac* transposon. Deletions of DMRs in the *Gtl2* and *H19* imprinted gene loci were engineered by simultaneous transfection with four expression vectors encoding CRISPR-Cas9 nucleases. (B) Sequences of PCR fragments amplified over the deleted regions confirmed the loss of both DMRs in DKO-phaESC-1 and DKO-phaESC-2. (C) Morphology of DKO-phaESC lines. Scale bar, 200 µm. (D) Haploid karyotypes were observed in both DKO-phaESC-1 and DKO-phaESC-2. (E) Transcription of imprinted genes *Gtl2* and *H19*, both of which are maternally expressed and regulated by the *IG*- and *H19*-DMRs, was reduced in DKO-phaESC-1 and DKO-phaESC-2. Gene expression was normalized to *Gapdh* relative to the parental cell line. Data represents relative expression of each sample with the mean values and standard deviation (n = 2). ** P < 0.01; * P < 0.05; ns, non-significant.

The maternally expressed *Gtl2* gene maps to a large imprinted cluster on mouse chromosome 12 and is regulated by the paternally methylated *IG*-DMR [24]. The maternally expressed gene *H19* maps to the paternally expressed *Igf2* gene on chromosome 7 and is regulated by a shared *H19*-DMR. As expected, transcription of *Gtl2* and *H19* was lost in both DKO-phaESC-1 and DKO-phaESC-2 (Fig 1E). In addition, expression of the paternally expressed *Dlk1* and *Igf2* genes, which are adjacent to *Gtl2* and *H19*, respectively, were slightly reduced in DKO-phaESC-1 and DKO-phaESC-2 compared with the expression of the parental phaESC line.

### Generation of semi-cloned embryos and scESCs by injection of DKO-phaESCs into oocytes

For assessing the potential of DKO-phaESCs as sperm replacement, we injected single cells into MII oocytes that were obtained from superovulated B6D2F1 female mice (Fig 2A). Previously, ahaESCs that were arrested in mitosis at M-phase had been used as sperm substitute with greater efficiency than ahaESCs in G0- or G1-phase, whereby the extrusion of the second and a pseudo polar body after injection were observed [9]. Following this report, we treated DKO-phaESC-2 with demecolcine and purified the metaphase arrested population of haploid cells by cell sorting (Fig 2B).

**Fig 2.**
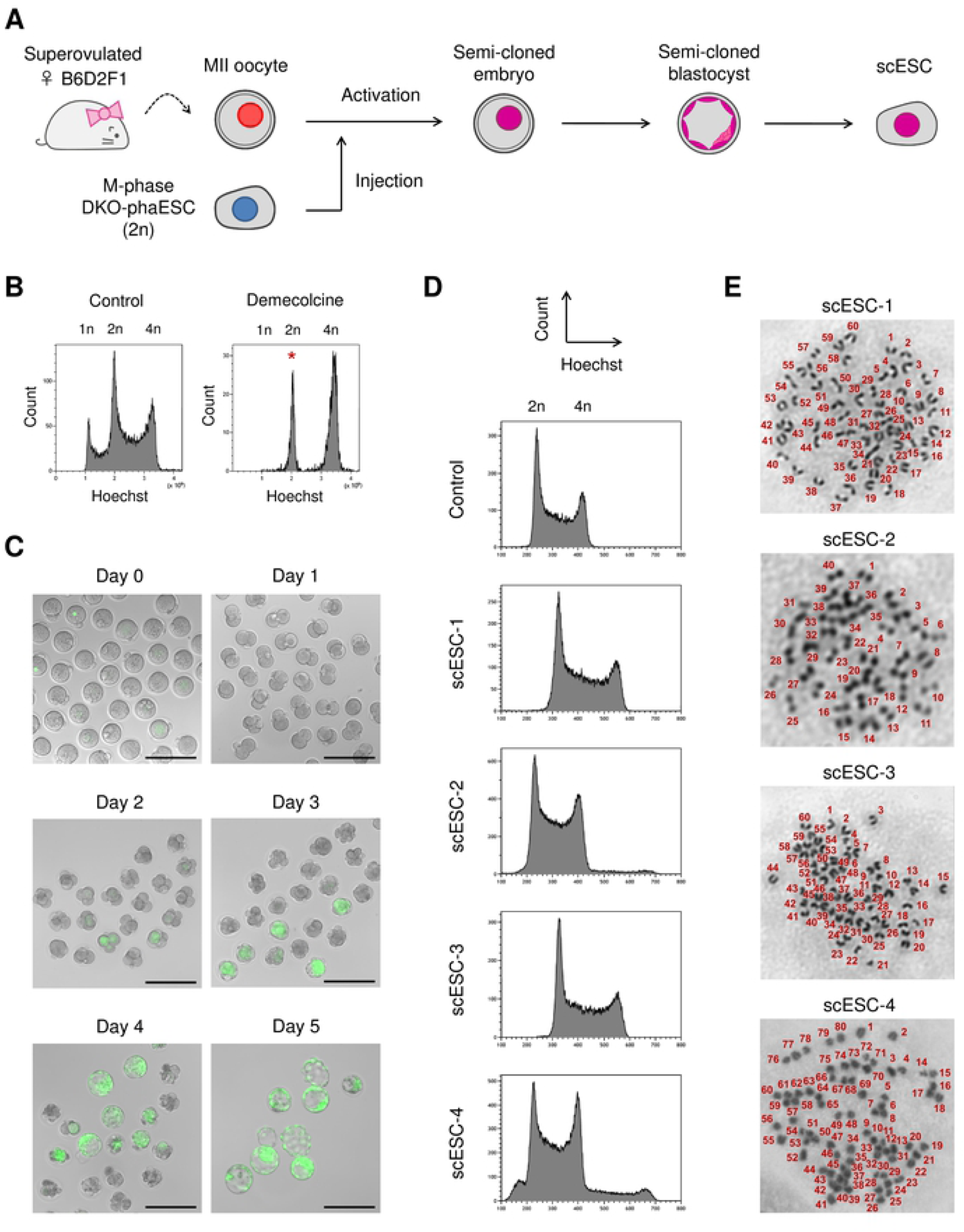
Characterization of scESC lines derived by injection of DKO-phaESCs into oocytes. (A) A scheme of the generation of scESC lines by injection of DKO-phaESCs into MII oocytes. (B) DKO-phaESCs were arrested in metaphase with demecolcine for 8 hours and sorted for a 2n DNA. The peak of the Hoechst intensity corresponding to 2n DKO-phaESCs is indicated (asterisk). (C) Semi-cloned embryo development after injection of DKO-phaESCs into oocytes. EGFP fluorescence merged with bright field images are shown. At day 4, morulae developed to blastocysts. Scale bar, 200 µm. (D) DNA content analysis of 4 scESC lines using flow cytometer after Hoechst staining. DNA content at the G1 phase of scESC-1 and scESC-3 appeared in the middle between the DNA content of G1 and G2 phase control diploid ESCs, indicating scESC-1 and scESC-3 are triploid. scESC-4 contained both diploid and tetraploid cells. (E) Metaphase spreads showed triploid karyotypes of scESC-1 and scESC-3, and a tetraploid karyotype of scESC-4.

Semi-cloned embryos were constructed by injection of M-phase DKO-phaESC-2 cells into MII oocytes, followed by activation with strontium chloride. After activation, the majority of semi-cloned embryos exhibited weak EGFP fluorescence distributed over the cytoplasm. However, a small number of embryos displayed a single round area of intense EGFP expression (Fig 2C), which indicates that the plasma membrane of DKO-phaESC-2 cells remained intact during injection. Semi-cloned embryos were cultured *in vitro* and developed to the 2-cell stage at the ratio of 58.7% (98/167), whereby little or no EGFP expression was detected (Table 1). Four-cell embryos initiated EGFP expression at day 2 after injection, followed by development to blastocysts at the ratio of 12.6% (21/167). All the blastocysts exhibited EGFP expression, indicating DKO-phaESC-2 cells contributed to the blastocyst genome and no parthenogenetic blastocysts developed.

**Table 1.** Summary of preimplantation development of semi-cloned embryos derived by injection of DKO-phaESCs into oocytes.

| No. of oocytes injected | No. of 2-cell embryos | No. of 4-cell embryos | No. of morulae | No. of blastocysts (% of oocytes injected) |
| --- | --- | --- | --- | --- |
| 167 | 98 | 50 | 43 | 21 (12.6%) |

To further analyze the semi-cloned blastocysts, we established 4 semi-cloned ESC lines scESC-1 to scESC-4 (S2A Fig). Genotyping revealed that these scESC lines possessed both wild-type and deleted alleles of the *IG*-DMR and *H19*-DMR, confirming the contribution of the DKO-phaESC and the oocyte genomes (S2B Fig). We next analysed the DNA content of all 4 scESC lines by Hoechst staining and flow cytometry (Fig 2D). scESC-2 exhibited an expected diploid DNA content, while the other 3 scESC lines appeared to be polyploid. Two scESC lines, scESC-1 and scESC-3, had a triploid DNA content, and scESC-4 contained cells with a diploid and tetraploid DNA content. The analysis of metaphase chromosomes confirmed a triploid karyotype in scESC-1 and scESC-3, and a tetraploid karyotype in scESC-4 (Fig 2E). The observation of polyploidy in 3 out of 4 scESC lines suggests that abnormal chromosome segregation in semi-cloned embryos is a frequent event. Considering that polyploidy is not compatible with mouse development [21, 22], it might be a key factor limiting the yield of semi-cloned mice.

### Generation of semi-cloned mice from semi-cloned embryos

In order to investigate whether semi-cloned embryos are competent to develop to mice, we injected DKO-phaESC-2 cells into MII oocytes that were obtained from superovulated B6D2F1 females. Subsequently, semi-cloned embryos were cultured to the 2-cell stage and transferred to oviducts of pseudopregnant Swiss Webster females. We chose Swiss Webster recipients for their albino coat color, which is readily distinguished from the agouti coat color of phaESCs and B6D2F1 oocytes. In parallel, albino 2-cell embryos were derived from Swiss Webster mice by *in vitro* fertilization (IVF) as a technical control. A total of 39 semi-cloned and 20 control 2-cell embryos were transferred to 4 recipient females (Table 2). Two of the four recipient females maintained pregnancy and delivered 6 pups (termed progeny no.1-6) and 1 pup (progeny no.7) (Fig 3A). Progeny no. 6 and 7 had dark eye color and toe biopsies indicated EGFP expression under UV illumination (Fig 3B). Genotyping confirmed that progeny no. 6 and 7 were female and heterozygous for the *IG*-DMR and *H19*-DMR, carrying wild-type and deletion alleles (Fig 3C). Both mice grew normally without any apparent phenotypes or health problems and delivered full-term pups after mating with Swiss Webster males (Fig 3D). Transmission of EGFP transgene was observed in about half of these pups (7/15) in the expected Mendelian manner (S3 Fig). Bisulfite DNA sequencing demonstrated that progeny no. 6 and 7 carried both methylated and unmethylated DMR alleles in 3 imprinted genes, *Kcnq1, Igf2r* and *Peg13* (Fig 3E). Some methylation patterns including *Kcnq1* and *Peg13* of progeny no.6 and *Igf2r* of progeny no. 7 appeared slightly hypermethylated. Considering the similarity to the control mouse, the 2 semi-cloned mice possessed normal methylation patterns in the 3 imprinted genes that we investigated.

**Table 2.** Summary of semi-cloned mice generation by injection of DKO-phaESCs into oocytes.

| Embryo types | No. of oocytes injected | No. of 2-cell embryos | No. of transferred 2-cell embryos | No. of delivered pups (% of transferred 2-cell embryos) |
| --- | --- | --- | --- | --- |
| Control | - | - | 20 | 5 (25%) |
| Semi-cloned | 50 | 39 | 39 | 2 (5.1%) |

**Fig 3.**
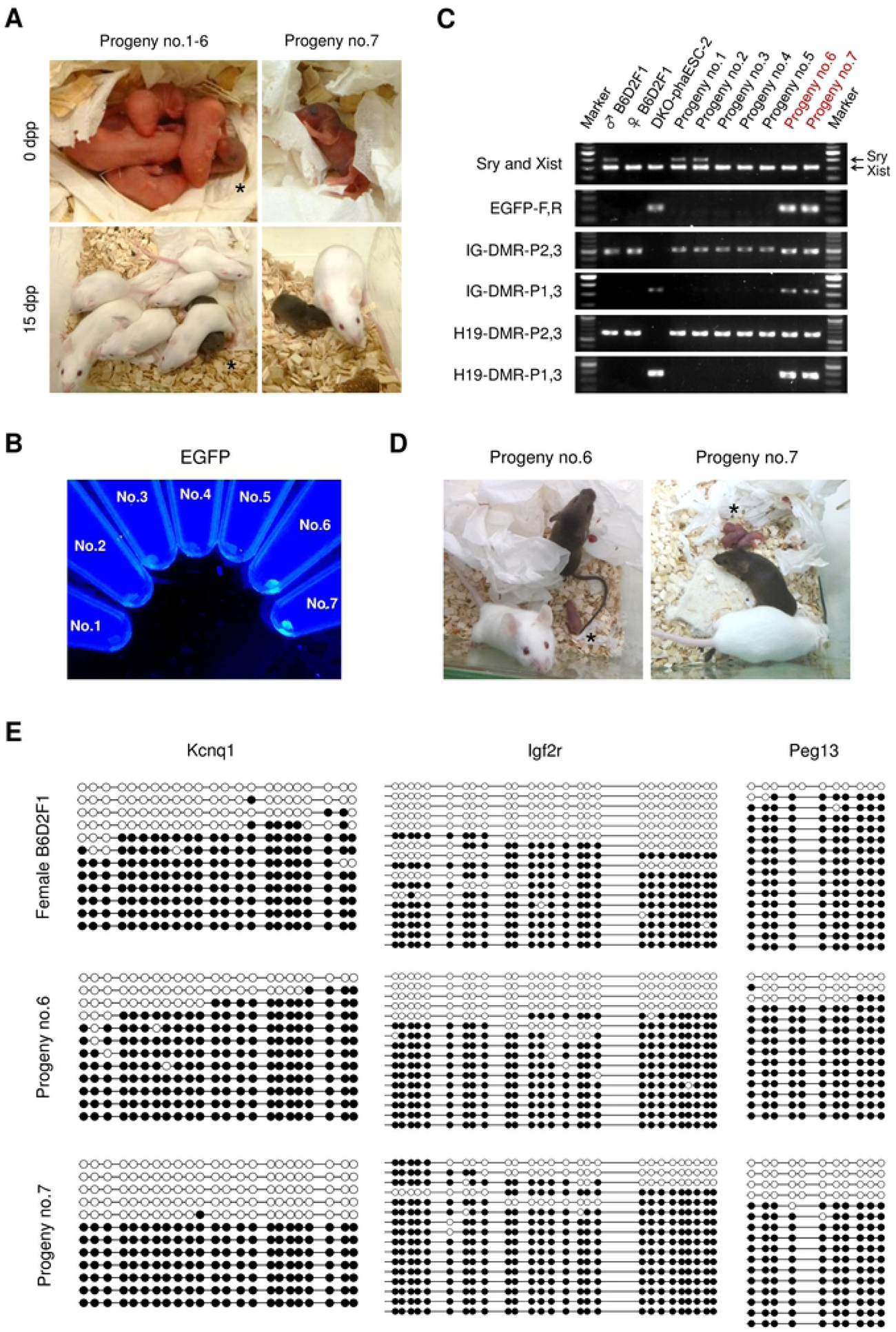
Generation of semi-cloned mice by transfer of semi-cloned embryos into recipient mothers. (A) 7 offspring (progeny no. 1-7) were obtained from 2 albino recipient mothers after transfer of semi-cloned 2-cell embryos. Progeny no. 6 (indicated by asterisk) and no. 7 displayed black eyes and agouti coat color indicating DKO-phaESC derived pigmentation. (B) Toe biopsies of progeny no. 1-7. Biopsies of progeny no. 6 and 7 expressed EGFP under UV illumination. (C) Genotyping of progeny. Progeny no. 6 and 7 possessed wild type and deletion alleles of the *IG*-DMR and *H19*-DMR. (D) Mating of semi-cloned progeny no. 6 and 7 with wild type males yielded healthy pups (indicated by asterisk). (E) Bisulfite DNA methylation analysis of *Kcnq1, Igf2r* and *Peg13* in biopsies of semi-cloned and control mice. White circles represent unmethylated CpGs; black circles represent methylated CpGs.

## Discussion

The successful production of fertile mice demonstrated that phaESCs with deletions of the *IG*-DMR and *H19*-DMR can be used as sperm replacement for mouse development. We used DKO-phaESCs for injection in this study, that had been treated with demecolcine to harvest 2 sets of chromosomes at the onset of mitotic spindle formation in M-phase. According to the previous study, M-phase ahaESCs contributed to semi-cloned embryos better than G0- or G1-phase ahaESCs [9]. The study reported the extrusion of the second polar body and pseudo polar body after the injection of M-phase ahaESCs. Consistent with this report, we observed the extrusion of two polar bodies after the injection of DKO-phaESCs in our study (Fig 2C). Semi-cloned blastocysts and semi-cloned mice were efficiently generated at the ratio of 12.6% against oocytes injected (21/167) and 5.1% against transferred 2-cell embryos (2/39), respectively (Table 1,2). The results indicate that the cell cycle between DKO-phaESCs arrested at M-phase and oocytes arrested at MII were well synchronized.

Meanwhile, we unexpectedly observed that some scESC lines derived from semi-cloned embryos were triploid or contained a mixture of diploid and tetraploid cells (Fig 2D and E). Several reasons can be assumed why these polyploid cells developed. For example, it is possible that diploidization of DKO-phaESCs before the first cleavage of semi-cloned embryos occurred or diploid ESCs were erroneously injected. In these cases, 2n chromatids of DKO-phaESCs would contribute to an embryo, resulting in the generation of a triploid embryo. A mixed karyotype of diploidy and tetraploidy was also observed in a scESC line. This is presumably caused by erroneous chromosome segregation at the 2nd or later cleavage [25]. A consequence of segregation defect possibly caused tetraploidy in a blastomere, while other blastomere(s) might have proceeded normal chromosome segregation and maintained diploid karyotype. As a result, both diploid and tetraploid cells might have developed in an embryo. Several studies on chimeric blastocysts containing diploid and tetraploid embryonic cells have shown that tetraploid cells contributed to extra-embryonic tissues but rarely to the fetus [26-28]. Nevertheless, homogenously tetraploid embryos formed blastocysts similar to diploid embryos during preimplantation development although their development was retarded before 15 days of gestation [21, 22]. Considering these results, it is reasonable that tetraploid cells developed to inner cell mass of the blastocyst and a scESC line containing diploid and tetraploid cells was generated in our study. Various possibilities can be considered as the cause of polyploidy by haESC injection. Unexpected polyploidy is possibly an impediment for the development of semi-cloned mice and could be a target for improvement to increase the yield of normal semi-cloned embryos in future. Further studies are expected to reveal the mechanism of polyploidy in semi-cloned embryos.

The uniqueness of haESC as replacement of gametic genome possesses substantial potential of applications especially because genetic mutations can be efficiently introduced into haESCs in contrast to oocytes and spermatozoa. We demonstrated that manipulation of haESCs allowed for simultaneous deletion of 2 DMRs in a single step with maintenance of a haploid karyotype. The generation of transgenic mice is one considerable application of haESCs as gametic genome replacement. Our data confirms previous reports that haESCs were suitable for generating semi-cloned embryos with high efficiency. Also, haESCs can be a powerful tool for genetic screening of factors required for fertilization or embryogenesis through injection of genetically modified haESCs into oocytes. To date, remarkable studies have reported the application of this haESC technology for genetic screening [16-18]. The mechanism how haESCs contribute to semi-cloned embryos remains an important focus for further study. Unexpectedly, polyploidy of semi-cloned embryos was frequent in our study. Further mechanistic insight on haESC contribution to embryos are needed for better understanding of gametic genome adaptation, which could also help to increase the efficiency of semi-cloning.

## Materials and Methods

### Animals and experiments

C57BL/6J and DBA/2J mice were purchased from Charles River Laboratories (Wilmington, USA). Swiss Webster and 129S6/SvEvTac mice were purchased from Taconic Biosciences (Rensselaer, USA). All the mice were housed in the animal facility of ETH Zurich. All animal experiments were performed under the license ZH152/17 in accordance to the standards and regulations of the Cantonal Ethics Commission Zurich.

### Oocyte collection

Four- to five-week-old female mice were induced to superovulate by injection of 5 IU pregnant mare’s serum gonadotropin followed by 5 IU human chorionic gonadotropin (hCG). Cumulus-oocyte complexes (COCs) were collected from the oviducts 15-17 hours after hCG injection and were placed in M2 medium. COCs were treated with 0.1% hyaluronidase until the cumulus cells disperses as indicated.

### Derivation and culture of phaESC lines

Derivation of phaESC lines from 129S6/SvEvTac mice was performed as previously described [7]. For introducing deletions of the *IG*-DMR and *H19*-DMR using the CRISPR-Cas9 system, previously published oligonucleotides for guide RNAs (gRNAs) [16] were ligated into the pX330-U6-Chimeric_BB-CBh-hSpCas9 vector (Addgene, #42230) that was digested with BbsI restriction enzyme (Fig. S1). Sequences of gRNAs are listed in Table S1. Simultaneous transfection of 4 Cas9/gRNA vectors, a *piggyBac* plasmid carrying a CAG-EGFP-IRES-hygro transgene, and a hyperactive *piggyBac* transposase plasmid was performed into a phaESC line using lipofectamine 2000 by following a manufacture’s protocol. Subsequently single EGFP expressing haploid cells were isolated by flow cytometer (MoFlo Astrios EQ, Beckman Coulter) after staining with 15 µg/ml Hoechst 33342 (Invitrogen). After the growth of clonal single colonies, a subset of cells in each line were stained with Hoechst and analyzed by flow cytometer to select cell lines containing haploid cells. Each haploid cell line was maintained without mouse embryonic fibroblasts and were subjected to genotyping and karyotyping. Purification of haploid 1n cell population in each phaESC line was performed by cell sorting after staining with Hoechst every 4-6 passages.

### Construction of semi-cloned embryos

Construction of semi-cloned embryos was performed following a published protocol [16] with a few modifications. Parthenogenetic haESCs with deletions of the *IG*-DMR and *H19*-DMR were arrested at M-phase by culturing in medium containing 0.05 mg/ml demecolcine (Merck) for 8 hours. After staining with Hoechst, DKO-phaESCs with a 2n DNA content were sorted by flow cytometer. In parallel, MII oocytes were harvested from superovulated B6D2F1 females. To construct semi-cloned embryos, sorted single cells were injected into MII oocytes using a piezo-driven micromanipulator (Eclipse Ti, Nikon; PiezoXpert, Eppendorf). After injection embryos were cultured in M16 medium for 1 hour and subsequently activated for 6 hours in KSOM medium containing 5 mM strontium chloride and 2 mM EGTA. After activation, embryos were washed and cultured in KSOM medium at 37°C under 5% CO_2_ in air.

### Genotyping

DNA extraction from cells and biopsies was performed using lysis buffer (100 mM Tris pH 8.5, 200 mM NaCl, 5 mM EDTA and 0.2% SDS) supplemented with 0.1 mg/ml proteinase K at 55°C for at least 4 hours. Debris were pelleted by centrifuging for 5 minutes at 13,000 rpm. Supernatant was replaced into a new tube containing equal volume of isopropanol. After mixing, the tube was centrifuged for 5 minutes at 13,000 rpm to pellet precipitated genomic DNA. The pellet was washed with 70% ethanol and resuspended by 50-200 μl water. PCR was performed using Phusion Hot Start II DNA Polymerase (Thermo Fisher Scientific) following the manufacturer’s protocol. PCR products were separated by electrophoresis on 1.5% agarose gels and stained with ethidium bromide for visualization under a UV transilluminator. Primers used for genotyping are listed in Table S1.

### Transcription analysis

RNA was extracted using the RNeasy Mini Kit (Qiagen) following the manufacturer’s protocol, including an on-column DNA digest using RNase-free DNase (Qiagen). RNA concentration was determined using a NanoDrop Lite (Thermo Fisher Scientific). 500 ng total RNA was reverse transcribed using the PrimeScript RT Master Mix (Takara) according to the manufacturer’s instruction. RT-PCR was performed at a 384 well format on the 480 Lightcycler instrument (Roche) using KAPA SYBR FAST qPCR KIT (Kapa Biosystems). Fold change expression was calculated using the ΔΔct method. *Gapdh* expression was used for normalization. Primers used for transcription analysis are listed in Table S1.

### Chromosome counting

For karyotyping, chromosome spreads of ESCs were prepared on glass slides as described [29]. Chromosomes were stained with Giemsa solution (Merck), washed with Gurr’s buffer, and subsequently chromosomes were imaged under the microscope (Axio Observer Z1, Zeiss). Pictures were taken using an ORCA-Flash4.0 camera (Hamamatsu Photonics K.K.) and chromosome counts were determined.

### *In vitro* fertilization (IVF)

Sperm mass collected from the cauda epididymis of Swiss Webster males were pre-incubated in Sequential Fert (ORIGIO) at 37°C under 5% CO_2_ in air. COCs were harvested from the oviductal ampulla of superovulated Swiss Webster females. After 1 hour of pre-incubation of sperm mass, a small aliquot of sperm suspension was added to a Sequential Fert drop containing COCs. Six hours later, oocytes were washed and transferred to KSOM medium. Embryo development to the 2-cell stage was assessed after 24 hours of IVF.

### Embryo transfer

Recipient Swiss Webster females were mated with vasectomized Swiss Webster males the night before, and plugs were confirmed in the morning of the day of the embryo transfer. Nine or ten 2-cell embryos derived by DKO-phaESC injection and 5 control 2-cell embryos by IVF were transferred into the oviducts of pseudo-pregnant recipient females. On day 19.5 of gestation, full-term pups were naturally delivered from recipient females.

### Bisulfite sequencing

Genomic DNA was extracted from toes of newborn semi-cloned mice and ear biopsy of 3 weeks old B6D2F1 mice with lysis buffer containing proteinase K, followed by isopropanol precipitation. Bisulfite conversion was performed using the EZ DNA methylation Gold kit (ZYMO Research). PCR was performed under the following temperature profile: 30 sec 98°C, 20 × (10 sec 98°C, 30 sec 65-55°C with −0.5°C per cycle, 30 sec 72°C), 35 × (10 sec 98°C, 30 sec 55°C, 30 sec 72°C), 5 min 72°C. The PCR products were cloned into pJet1.2 vector using the CloneJET PCR Cloning Kit (Thermo Fisher Scientific), followed by the transformation into competent DH5α *E*.*coli*. Insert sequences for each colony were obtained through the commercial Ecoli NightSeq service (Microsynth). Bisulfite sequencing was analyzed with the QUMA methylation analysis tool (http://quma.cdb.riken.jp/). Primers used for PCR and sequencing are listed in Table S1.

### Statistical analysis

For comparison of quantitative RNA expression levels of imprinted genes, measurements were analyzed with the GraphPad Prism 8 software using a two-tailed unpaired t-test. A p-value < 0.05 was considered statistically significant.

## Acknowledgments

We thank Mr. Stefan Butz and Dr. Tuncay Baubec for providing primers and advice on bisulfite sequencing. We also acknowledge Ms. Michèle Schaffner and Mr. Thomas M. Hennek for their technical support on embryo transfer. This work was supported by the Swiss National Science Foundation (grant 31003A_152814/1).

## Supporting Information

**S1 Fig. Deletions of the *IG*-DMR and *H19*-DMR in phaESC lines**. (A) A design of gRNAs and primers targeting the deletions of the *IG*-DMR. (B) A design of gRNAs and primers targeting the deletions of the *H19*-DMR. (C) PCR fragments flanking both *IG*-DMR (319 bp) and *H19*-DMR (407 bp) by primers targeting deleted loci were observed in 2 DKO-phaESC lines, whereas the deleted sequences were absent in DKO-phaESC-1 and DKO-phaESC-2.

**S2 Fig. Derivation and genotyping of scESC lines**. (A) Derivation of scESC lines from blastocysts generated by injection of DKO-phaESCs into oocytes. Images of blastocysts, outgrowth (passage 0) and scESCs after derivation are shown. Regular black bar, 100 µm; bold black bar, 200 µm; white bar, 100 µm. (B) Genotyping of 3 scESC lines. All 3 scESC lines exhibited both wild type and mutant alleles for the *IG*-DMR and *H19*-DMR, indicating both oocytes and DKO-phaESCs genome contributed to the genome of blastocysts.

**S3 Fig. Genotyping of pups born to semi-cloned mice**. PCR-based genotyping was performed for 15 pups born to semi-cloned females (progeny no.6 and 7) and wild type Swiss Webster males. EGFP transgene was inherited to 7 among 15 pups.

**S1 Table. List of oligos**.

